# Trapline foraging by nectar-collecting hornets

**DOI:** 10.1101/2024.12.22.629945

**Authors:** Mathilde Lacombrade, Kristine Abenis, Charlotte Doussot, Loïc Goulefert, Kenji Nanba, Jean-Marc Bonzom, Mathieu Lihoreau

**Author notes:** These authors contributed equally to the work (Mathilde Lacombrade, Kristine Abenis).

## Abstract

Central place foraging bees, butterflies, birds and bats are known to develop routes in order to visit familiar plant resources in a stable and repeatable order called “traplines”. Here we report a similar behaviour in a social wasp, the Japanese yellow hornet *Vespa simillima*. We monitored the foraging movements of individually marked wild hornets collecting sucrose solution on four artificial flowers placed in the field. After thirty foraging bouts, all the hornets had developed a repeatable flower visitation sequence. Using two different arrays of flowers, we show how hornets increase their foraging efficiency through time but do not always use the shortest path to visit all flowers, often favoring movements between nearest neighbor flowers over global path optimization. Our study adds nectar-foraging wasps to the growing list of animals developing traplines thereby opening new perspectives for comparative cognition research.

## Introduction

Nectar-feeding animals must typically visit dozens to thousands of flowers within a single foraging trip in order to fill their crop to capacity. In central place foragers, such as bees, hummingbirds, and many bats, individual foragers tend to use repeatable foraging circuits to exploit known food locations that renew over time (Ohashi et al. 2009; Lihoreau et al. 2013; Tello-Ramos et al. 2015). This routing behaviour, known as “trapline foraging” (Thomson et al. 1997), provides foragers substantial benefits in terms of reduced competition and increased rate of energy gain (Possingham 1989).

The development of traplines has been described in details by observing animals exploiting feeders (i.e. artificial flowers) over several consecutive hours. In this approach, bumblebees and honey bees foraging alone in an array of four to ten artificial flowers in which nectar load was renewed at high rates, frequently use the shortest route to visit all flowers once and return to their nest (Buatois et al. 2024). Through trial and error, foragers tend to reduce their revisits to empty flowers, use straight flight paths between visiting flowers, and arrange their flower visitation sequences in such a way that minimize overall travel distances (Lihoreau et al. 2012a; Reynolds et al. 2013). Once established, a trapline can last for days or weeks provided that food resources remain stable and continue to replenish (Thomson et al. 1996).

In bees, route optimization is dependent on spatial scales. Foragers are more efficient at finding efficient traplines minimizing travel distances at large spatial scales when feeding locations are spaced by dozen to hundred meters (i.e. between flower patches) than at small spatial scales when locations are spaced by only a few centimeters or meters (i.e. within a flower patch) (Lihoreau et al. 2010; Lihoreau et al. 2012b). Presumably, in the latter case the energy cost of using a long, suboptimal, route is negligible compared to the cognitive cost involved in route optimization (Buatois and Lihoreau 2016). At both spatial scales, however, travel optimization is generally imperfect as bees tend to favour straight movements over sharp turns (Ohashi et al. 2007, Woodgate et al. 2017), and jumps between nearest flowers over longer strategic bypasses (Saleh et al. 2006).

Here we explored whether the development of such multi-destination routes, previously described in several central place foraging nectarivore species can also be observed in wasps. Just like bees, social wasps must collect large quantities of floral nectar to provision their nests (Bouchebti et al. 2022). Previous studies using mark-recapture approaches have showed that Asian yellow-legged hornets (*Vespa velutina*) often return to the same foraging places (Monceau et al. 2014; Ueno 2015), thus setting the stage for place memories and an ability to use traplines in hornets. In the current study, we tested this ability in a population of wild Japanese yellow hornets *Vespa simillima* foraging in two arrays of four artificial flowers in the field.

## Methods

### Study site and insects

The experiments took place on a study site located in Minamisoma city (Fukushima prefecture, Japan: 37°37’35.612’’N, 140°54’17.567’’E) in September 2023. We worked on a private road bordered by cherry trees on which six honey bee hives (*Apis mellifera*) had been placed in May 2023. A population of wild yellow hornets (*V. simillima*) was predating on the honey bees. We attracted the hornets by placing a gravity feeder containing *ad libitum* 40% (v/v) sucrose solution in the middle of the experimental site (see pre-training location marked with a pink disc in Fig. 1). When hornets landed on the feeder, we paint-marked them with enamel paint on the thorax, the abdomen, or both, for individual recognition. After collecting sucrose solution, the marked hornets flew away in straight lines towards their colony nests. Detailed observations of these return flights indicated that there were three different nests in the study site (see arrows in Fig. 1).

**Fig 1.**
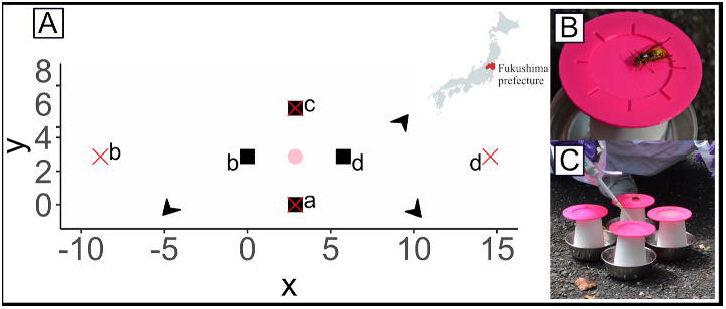
Arrays of artificial flowers. A) We settled artificial flowers (a-d) on a road in the middle of our experimental site in the Fukushima prefecture, Japan (37°37’35.612’’N, 140°54’17.567’’E, map produced with MapChart). The pink disk highlights the pretraining location. The small array of flowers is indicated with black squares - flower coordinates (x, y): a(2.875, 0), b(0, 2.875), c(2.875, 5.750), d(5.750, 2.875). The large array of flowers is indicated with red crosses - flower coordinates: a (2.875, 0), b(-8.850,2.875), c(2.875, 5.750), d(14.600, 2.875). Black arrows indicate the directions of hornet nests (precise locations are unknown). B) Picture of a paint-marked hornet collecting sucrose solution on an artificial flower. C) Picture of the patch of four flowers used for pre-training.

### Pre-training

Once we had identified several hornet foragers frequently returning to the feeder, we caught them all in individual cages in order to avoid aggressive interactions between hornets on feeders that could interfere with their natural foraging behaviour (personal observations). One hornet (i.e. the focal individual) was then released for testing. This hornet was presented four identical feeders (hereafter called “artificial flowers”) arranged in a patch at the pre-training location directly on the ground (flowers were interspaced by 5 cm; Fig. 1A). Each flower was made of a large 3D-printed pink plastic landing platform (diameter 10cm) placed on top of a transparent cylindrical paper cup (diameter 5cm, height 8cm; Fig. 1B). In the middle of the landing platform, we provided a controlled volume of 40% sucrose solution with a micropipette (Genex beta P20). The flower was placed in a metallic cup (diameter 12cm, height 3cm) filled with water in order to avoid that ants could climb to the platform and feed the sucrose reward.

We measured the nectar crop capacity of the focal hornet (i.e. maximum amount of liquid the hornet can collect to fill its crop to capacity) by presenting drops of 10uL 40% sucrose solution on each of the four flowers and refilling them with a new drop of the same volume each time the focal hornet collected them (Lihoreau et al. 2010) (Fig. 1C). We monitored the total volume of nectar collected by the hornet during a single foraging bout (i.e. foraging sequence starting when the hornet came to visit the array of flowers and ending when the hornet flew back to its nest) over five consecutive foraging bouts. We then estimated the crop capacity of each hornet by averaging the volume collected during the five foraging bouts (min= 152ul, max=360ul, n = 7 hornets).

### Training

Once the crop capacity of the focal hornet was estimated and the hornet had returned to its nest, we moved the four artificial flowers into their final position (see flower coordinates in Fig. 1A). We used two different arrays of flowers. In the “small array”, the four flowers were arranged in a square of 4m side. In this array, a hornet moving between nearest neighbour flowers would use the shortest possible route to visit all flowers at once. In the “large array” the four flowers were arranged in a diamond of 23.45m x 5.75m side. In this array, a hornet moving between nearest neighbour flowers would use a long, suboptimal, route (i.e. the optimal route would be to circulate around the array).

For both arrays, at the beginning of a foraging bout, each flower contained 1/4th of the nectar crop capacity of the focal hornet, so that the insect had to visit the four flowers in order to fill its crop to capacity and return to its nest. Each focal hornet was observed foraging in a given array for 30 consecutive bouts (average 113.15 ± (sd) 26.50 minutes of observations per hornet, n = 7 hornets).

We collected the data using a custom-made computer program enabling us to record events (time of occurrence of a given behaviour) by pressing a key (Lihoreau et al. 2010). At each foraging bout of each hornet, we recorded the time at which the hornet landed and took off from each flower, as well as the time at which the hornet returned to its nest. In this approach, we tested 5 hornets in the small array and 3 hornets in the large array. Each hornet was tested on a different day. After testing a focal hornet, all the other captive hornets, previously trained and marked, were released for later tests.

### Data analysis

We performed all the statistical analyses with R, and produced figures and tables with R and Python. Flower visitation sequences by each hornet (excluding immediate revisits) are available in the Fig. S1. Before any analysis, we excluded from the raw data all revisits to flowers after the 4 flowers of the array have been visited within the same foraging bout (i.e. all flowers empty; Lihoreau et al. 2010). These revisits occurred in less than 5% of all foraging bouts. We then assessed the repeatability and the optimality of flower visitation sequences for each foraging bout of each hornet using four parameters: the determinism index (DET), the revisit rate, the crop filling rate, and the relative difference between travelled distance and optimal route (see details below).

The DET calculation follows methods described by Ayers et al. (2015) and Buatois and Lihoreau (2016). This index indicates how much an animal repeats sequences of visits between multiple locations through time: an index of 1 indicates a perfectly repeated sequence of visits, while an index of 0 indicates the complete absence of repetition. The DET takes into account sequence length and revisits as well. Here we ignored sequences that included less than three visits. Sequences were considered similar when they followed exactly the same order, i.e. the route linking flowers abcd was different from the route dcba. We grouped the foraging bouts of each hornet into bins of five in order to measure the DET, assess its evolution through time, and the impact of the flower array size.

We analyzed the tendency of hornets to revisit the same flower several times during a given foraging bout (i.e. unrewarded revisits). The rate of immediate revisits corresponded to the number of times a hornet returned to a flower without visiting any other flowers in between. The rate of non-immediate revisits corresponded to revisits to a flower after having visited other flowers.

To calculate the nectar crop filling rate, we assumed the hornet’s crop capacity to be 25% filled when one flower containing a reward was visited (precise volume of nectar collected is unknown). The crop filling rate was calculated as the number of flowers visited divided by the total time spent per foraging bout.

The relative difference between observed sequences used by hornets and the shortest possible sequence was calculated as the difference between the estimated travel distance (assuming hornets flew straight lines between flowers; Lihoreau et al. 2010) and the optimal distance (i.e. shortest route to visit all flowers once by flying straight lines between flowers) divided by the optimal distance multiplied by 100. This indicated the deviation from optimal. Deviation was zero % when the hornet uses the optimal route. We calculated the estimated and optimal travel distances by adding the distance between the approximated nest location and the first visited flowers, the distance between flowers, and the distance from the last visited flower to the nest. We estimated the nest location using hornets’ homing directions at the end of each foraging bout through visual observations. We approximated the nest positions by placing them in a theoretical circle whose center is in the middle of the flower arrays and has a 20m radius.

All previously described variable changes through time (foraging bout number) were statistically investigated using Generalized Linear models (GLM). Yet, due to the small sample size (5 and 3 hornets tested in each flower array) the random effects were not added as we possess less than 5 grouping levels and not enough observations (Gelman and Hill 2006). We removed outliers for the crop filling following the interquartile rule (we removed observations that fall below Q1 − 1.5 IQR or above Q3 + 1.5 IQR.).

To visualize hornet movements, we created spatial movement networks (Fig. 2) illustrating all the transitions between artificial flowers by hornets using NetworkX Python (Hagberg et al. 2008). For a given hornet, these networks represent the frequency and direction of all movements between two flowers (Pasquaretta et al. 2019).

**Fig 2.**
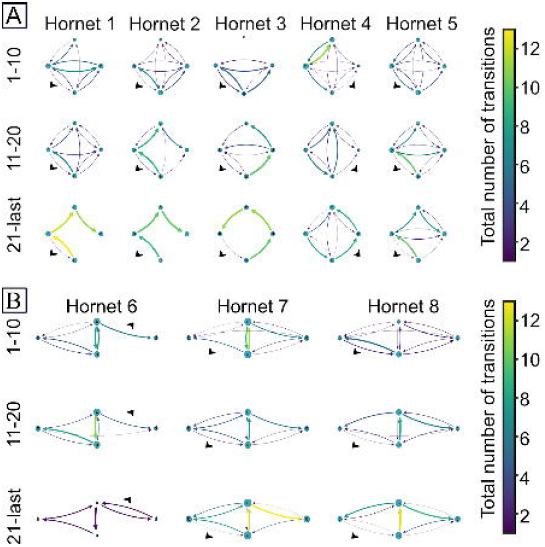
Spatial movement networks of hornets A) in the small array of flowers and B) in the large array of flowers. Columns show data for individual hornets. Rows represent bins of 10 foraging bouts (not all hornets completed 30 foraging bouts, see details in Fig. S1). Blue nodes represent flowers (see details about flower names and coordinates in Fig. 1A). The diameter of each node is proportional to the number of visits on those flowers for each bin of 10 foraging bouts. The thickness of each arrow is proportional to the total number of executed transitions for each bin of 10 foraging bouts. The direction of the arrows indicates the direction of the movement between flowers. The absolute number of transitions is indicated by the colormap. Black arrows indicate the directions of the nest of each hornet.

## Results

### Small flower array

We first tested whether hornets could develop traplines in a small array in which all flowers were regularly arranged and close to each other. In this array, the hornets used increasingly repeatable flowers visitation through time, as illustrated by the spatial movement networks in which arrows tend to be less numerous, brighter and larger in the last bin of foraging bouts than in the first bin (Fig. 2A). All five hornets increased their route repeatability through time (DET: GLM with gamma log link function, df=43, t=7.405, p-value<0.001; Fig. 3A). With experience, they reduced their number of immediate and non-immediate revisits to empty flowers (GLM gamma distribution with an inverse link function, immediate revisits: df=228, t=5.755, p<0.001; non-immediate revisits: df = 221, t=4.35, p-value <0.001, Fig. 3B), as well as the difference between traveled distance and the shortest possible route (difference to optimal: GLM gamma distribution with an inverse link function, df=132, t=3.00, p-value <0.001; 2.49 +/-10.24% in the last 5 foraging bout; Fig. 3C). In doing so, they also increased their crop filling rate (GLM gaussian distribution, df=221, t=8.036, p<0.001, Fig. 3D). Thus, increased route repeatability resulted in higher foraging efficiency through time with faster foraging bouts, reduced estimated travel distances, and higher rate of crop filling per minute.

**Fig 3.**
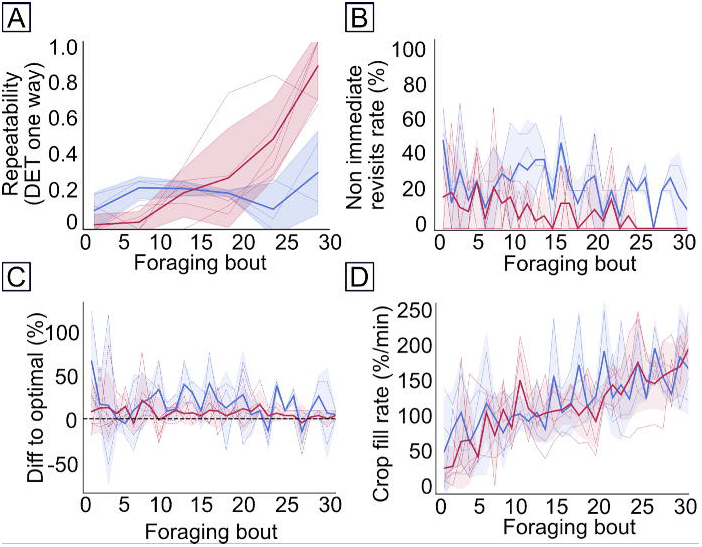
Foraging performances by hornets in the small (red, N=5 hornets) and large (blue, N = 3 hornets) array of flowers over 30 consecutive foraging bouts. (A) Route repeatability as measured by the determinism index (DET), B) Number of non-immediate revisits to empty flowers (in percentage), (C) Relative difference between traveled distance and optimal route minimizing travel distances (in percentage), D) nectar crop filling rate (in percentage per minute). Solid lines represent mean, colored areas the standard deviations, and colored dashed lines the individual data. In C, black dashed line represents the optimal route minimizing overall travel distance. Data under this line means individuals did not visit the 4 flowers.

Ultimately, the hornets used the shortest possible flower visitation sequence in 51.33% (SD = 19.63) of their foraging bouts, moving between nearest unvisited flowers in 76.4% of their transitions. Interestingly the first flower visited during the route and the direction in which the hornets turned in the array varied across hornets indicating that route establishment was the consequence of individual learning and memory (see Fig. S1). In particular, the hornets began their routes by visiting the flower that was the closest to their nest location in 87.5 (SD=4.61) % of the cases.

### Large flower array

We then tested whether hornets could still develop traplines in a larger array in which flowers were irregularly arranged and farther away from each other. In this array, the qualitative comparison of spatial movement networks again suggest that hornets used less numerous, brighter and larger transitions between flowers in the last bin than in the first bin of foraging bouts (Fig. 2B). The hornets did not significantly increase their route repeatability with experience (DET, GLM with gamma log link function, df=43, t=0.61, p-value=0.54; Fig. 3A). They also did not significantly reduce their number of non-immediate revisits through time (GLM with gamma distribution, non-immediate revisits: df=221, t=0.864, p-value=0.39; Fig. 3B). However, they increased their foraging efficiency as shown by the significant reduction of their number of immediate revisits to flowers (GLM with gamma distribution, immediate revisits: df=228, t=4.063, p<0.001), decrease of overall travel distance (difference to optimal: GLM gamma distribution with an inverse link function, df=132, t=2.59, p=0.01; 16.72 SD=23.13% in the last 5 foraging bout; Fig. 3C), and increase of nectar crop filling rate (crop fill rate: GLM, df=221, t=5.47, p-value<0.001, Fig. 3D) with experience.

In this large array, the hornets used the shortest possible sequence to visit all flowers in 30.38% (SD=11.25) of their foraging bouts (see detailed sequence visitation in Fig. S1), favoring movements between nearest unvisited neighbour flowers in 61.19% of their transitions. Here again the first flower visited during the route and the direction in which the hornets turned in the array varied across hornets, with hornets beginning their routes by visiting the flower the closest to their nest location in average 87.5% of the cases (see Fig. S1).

Overall, our results suggest poorer route development performances by hornets in the large array compared to smaller one (DET: df=43, t=-3.846, p-value<0.001; non-immediate revisits: df=221, t=3.69, p<0.001). However, the overall foraging efficiency, as measured by crop filling rate and relative traveled distance to the optimal route, was similar in both conditions (crop filling rate: df=221, t=-0.17, p-value=0.867, difference to optimal: df=132, t=1.09, p=0.278).

## Discussion

Nectar-feeding bees, butterflies, birds and bats learn to follow traplines to efficiently exploit plant resources in their environment (Ohashi and Thomson 2009; Lihoreau et al. 2013, Tello-Ramos et al. 2015). Here we found that a nectar-collecting wasp, the Japanese yellow hornet, also develops traplines to forage on artificial flowers, thereby comforting the idea that this routing behaviour is a widespread adaptation to central place foraging for nectar (Rombault et al. 2022).

Traplining by hornets was most evident in the small array of flowers, when flowers were arranged in a square of 4m side. In this condition all hornets gradually developed a route to visit all flowers once and return to their nest, either by turning right or left in the array. Route development was accompanied by a reduction of revisits to empty flowers, and a reduction of travel distances, thereby increasing foraging efficiency through time, and sometimes leading to the stabilization of the shortest possible route (see flower visitation sequences of hornets 1,2,3,5 in Fig. S1). This observation is very similar to those of bumblebees and honey bees collecting nectar on artificial flowers at similar spatial scales (Lihoreau et al. 2010, 2011; Buatois and Lihoreau 2016).

The hornets also reached high foraging efficiency in the larger flower array, when flowers were arranged in diamond of 12m side. In this context, they tended to make less revisits to empty flowers, increased their crop filling rate and reduced the difference between the distance they travelled and the optimal distance as they gained experience in the array. However, none of them stabilized a route. Relatively high levels of route repeatability were reached by favouring movements between nearest neighbour flowers, forcing them to detours from the shortest possible sequence (see visitation sequences of hornets 6,7,8 in Fig. S1). This observation is also coherent with trapline foraging studies in bumblebees indicating that bees tend to favor movement between nearest neighbours over sharp turns, even if this leads to suboptimal routes in flower arrays where nearest neighbour movements and travel optimization are put in conflict (Ohashi et al. 2007; Woodgate et al. 2017). Interestingly, and as expected from these studies on bees, hornets from different nests initiated their route by visits to the flower closest to their nest thereby confirming the importance of nest location in the final geometry of traplines.

The absence of travel optimization by hornets in our large array of flowers does not prove their inability to do so. Several reasons could explain this result. Firstly, it is possible that travel distances in our arrays were too small to incur significant movement costs, even in our large array, and trigger optimization behaviour. Bees, for instance, are more prone to optimizing their route geometry at large spatial scales, when feeding sites are spaced by several dozen meters and using long suboptimal routes is likely energetically costly (Lihoreau et al. 2012; Buatois and Lihoreau 2016). Yellow hornets have likely a similar (if not larger) foraging range as honey bees, as suggested by observations on similar sized yellow-legged hornets (*Vespa velutina)* which have been reported to forage up to at least five kilometers around their nest (Poidatz et al. 2018) and possess a maximum flight capacity of dozen kilometers (Sauvard et al. 2018). *Vespa*’s cuticles possess photovoltaic properties as it converts light into a metabolic form of flight energy (Ishay 2004) suggesting they have long distance flight capacities. The small difference between the longest and the shortest route in our large experimental array of flowers could therefore be negligible according to the flight costs and capacities of hornets. It is also possible that our observations were not long enough (30 trials, range: 76 min 47s - 158 min 5s) in order to enable the hornets to find and learn the shortest possible route in our setups. Previous studies with bumblebees show that the number of trials necessary for foragers to develop and optimize a route varies greatly with spatial scale and arrangement of artificial flowers (e.g. 20 trials in Lihoreau et al. 2010, 80 trials in Lihoreau et al. 2011). Future experiments should therefore explore the spatial foraging patterns of hornets in a wider range of spatial scales, numbers of feeding sites and training durations.

Our observation of trapline foraging in hornets indicates that these insects use spatial and visual memories when foraging for nectar, and highlight a strong resource fidelity of hornets. Future studies testing whether hornets, like bees, develop traplines between natural nectar resources and between the same flower species (flower constancy) will be necessary to further assess their contribution to pollination in a context where they also represent a threat for many pollinators (Brock et al. 2021; Rojas-Nossa et al. 2023).

## Acknowledgements

This work was funded by the Mission Interdisciplinaire CNRS-IRSN (MITI BEECONNECT) to JMB and ML, and the European Commission (ERC Cog BEEMOVE) to ML.

## Supplementary figures

**Fig S1.** Flower visitation sequences by hornets excluding immediate revisits. Coloured letters represent single visits to different flowers (see details about the spatial arrangements of flowers in Fig. 1A).

