## Supplementary figures and images for "Trapline foraging by nectar-collecting hornets"

### Fig S1

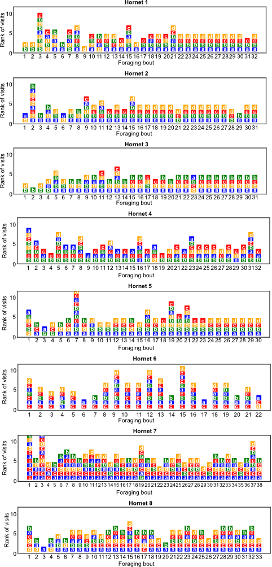
